# Loss of synaptopodin impairs mGluR5 and protein synthesis dependent mGluR-LTD at CA3-CA1 synapses

**DOI:** 10.1101/2023.08.02.551676

**Authors:** Pei You Wu, Linjia Ji, Claudia De Sanctis, Anna Francesconi, Yanis Inglebert, R. Anne McKinney

## Abstract

Metabotropic glutamate receptor-dependent long-term depression (mGluR-LTD) is an important form of synaptic plasticity that occurs in many regions of the CNS and is the underlying mechanism for several learning paradigms. In the hippocampus, mGluR-LTD is manifested by the weakening of synaptic transmission and elimination of dendritic spines. Interestingly, not all spines respond or undergo plasticity equally in response to mGluR-LTD. A subset of dendritic spines containing synaptopodin (SP), an actin-associated protein, are critical for mGluR-LTD and protect spines from elimination through mGluR1 activity. The precise cellular function of SP is still enigmatic and it is still unclear how SP contributes to the functional aspect of mGluR-LTD despite of its modulation on the structural plasticity. In the present study, we show that the lack of SP impairs mGluR-LTD by negatively affecting the mGluR5-dependent activity. Such impairment of mGluR5 activity is accompanied by a significant decrease of surface mGluR5 level in SP knockout (SPKO) mice. Intriguingly, the remaining mGluR-LTD becomes a protein synthesis-independent process in the SPKO and is mediated instead by endocannabinoid signaling. These data show for the first time that the postsynaptic protein SP can regulate the locus of expression of mGluR-LTD and provide insight to our understanding of spine/synapse-specific plasticity.

**Significance statement:** Hippocampal group I metabotropic glutamate receptor dependent long-term depression (mGluR-LTD), a form of learning and memory, is misregulated in many murine models of neurodevelopmental disorders. Despite extensive studies there is a paucity of information on the molecular mechanism underlying mGluR-LTD. Previously, we reported that loss of synaptopodin, an actin-associated protein found in a subset of mature dendritic spines, impairs mGluR-LTD. In the current study, we uncover the molecular and cellular deficits involved. We find that synaptopodin is required for the mGluR5-Homer interaction and uncover synaptopodin as a molecular switch for mGluR-LTD expression, as mGluR-LTD becomes protein synthesis-independent and relies on endocannabinoid signaling in synaptopodin knock-out. This work provides insight into synaptopodin as a gatekeeper to regulate mGluR-LTD at hippocampal synapses.

## Introduction

Synaptic plasticity, characterized by long-term potentiation (LTP) and long-term depression (LTD), is a major form of plasticity in brain that takes place at the level of individual synapses and is thought to be an underlying mechanism for learning and memory formation (1, 2). Dendritic spines, which make up the postsynaptic component of the synapse, are the primary loci of synaptic changes during plasticity. Dendritic spines can modify their molecular composition to alter synaptic activity and undergo long-term changes in their morphology based on the pattern and the strength of synaptic transmission (3). Specifically, LTD is known to induce a decrease in synaptic transmission, leading to the weakening or the elimination of synapses that are respectively associated with the shrinkage and the removal of the dendritic spines (4, 5). At hippocampal Schaffer collateral-CA1 (Sc-CA1) synapses, the most well studied form of LTD is NMDAR-LTD, which is typically induced by NMDAR activity through low frequency stimulation (4, 6–8). Another distinct form of LTD (mGluR-LTD) that requires the activation of group 1 metabotropic glutamate receptors, mGluR1 and mGluR5, was later discovered. It was found to be an important form of synaptic plasticity as it is altered in many neurological disorders such as Fragile X syndrome and Alzheimer’s Disease (9). Unlike NMDAR-LTD, hippocampal mGluR-LTD in mature animals depends on rapid *de novo* protein synthesis occurring in dendritic spines, although presynaptic mechanisms including retrograde endocannabinoid (eCB) signaling could also contribute to the plasticity (10–12). Interestingly, how dendritic spines respond to mGluR-LTD is controversial as some report spine loss during mGluR-LTD while others do not (13, 14). This differential expression of plasticity suggests that the spines are heterogeneous and opens up the question of how plasticity-responsive spines are different from the inert spines. Recently, studies on the postsynaptic protein synaptopodin (SP), an actin-associated protein, have begun providing insight onto spine-specific plasticity (15–17). SP is found specifically in a subset of mature dendritic spines and is responsible for the formation of the spine apparatus (SA), a specialized smooth endoplasmic reticulum (sER) derived organelle believed to serve as an intracellular Ca^2+^ reservoir in spines (17, 18). Although the exact function of SP in neurons remains largely enigmatic, it has been shown to play an indispensable role in different forms of synaptic plasticity and homeostatic plasticity (19–22). Previously, Holbro et al. has shown the spine-specific expression of mGluR-LTD that is dependent of the presence of sER in spines, suggesting perhaps the involvement of SP and SA (23). Subsequently, we demonstrated that SP is indeed involved in mGluR-LTD. We have shown that during mGluR-LTD, spines containing SP are protected from elimination or shrinkage, while the spines lacking SP are lost. Moreover, in SPKO mice, mGluR-LTD no longer induces spine loss, suggesting that SP is required for mGluR-LTD-induced structural plasticity (24). Although mGluR-LTD affects structural plasticity in a spine-specific and SP-dependent manner, it is unknown how mGluR-LTD-dependent functional plasticity is affected by SP in dendritic spines.

Here, to better understand the mechanisms underlying the mGluR-LTD deficit in SPKO at Sc-CA1 synapses, we assessed mGluR1 and mGluR5 activity individually in wild type (WT) and SPKO mice and found that mGluR5 activity is impaired in the SPKO. We found that mGluR5 surface expression is reduced in SPKO mice due to decreased mGluR5 association with Homer, a major scaffolding protein for mGluR1/5. Moreover, we report that mGluR-LTD becomes protein synthesis-independent and relies instead on endocannabinoid (eCB) signaling in the SPKO. These findings shed light on the function of SP in spines and during mGluR-LTD and reveal SP as a molecular switch that governs the mechanism of expression of mGluR-LTD.

## Results

### mGluR1 and mGluR5 differentially contribute to mGluR-LTD in absence of synaptopodin

At excitatory synapses, dendritic spines are the major locus for the expression of synaptic plasticity often manifested by alterations in both the functional and the structural properties of spines (3). Chemical activation of group 1 mGluRs with the agonist dihydroxyphenylglycine (DHPG) induces long-term synaptic depression at Schaffer collateral/Sc-CA1 synapses (mGluR-LTD) that is accompanied by the selective loss of spines that lack SP. To better understand the impact of SP on functional synaptic weakening, we induced LTD by brief application of DHPG (*S*-DHPG; 100 μM, 5 min) and recorded field potentials (fEPSPs) at Sc-CA1 synapses in acute slices from WT and SPKO mice (**Fig. 1A**). In WT, DHPG rapidly induced robust long-lasting depression of evoked fEPSPs measured 40 min after the end of DHPG application (59.1±4.8% of baseline, n=11; **Fig. 1A**). In contrast, in SPKO mice the DHPG-induced LTD was strongly weakened (76.5±4.4% of baseline, n=10; **Fig. 1B**, see **Suppl. Fig. 1** for WT vs. SPKO), in confirmation of our previous findings (24). Both mGluR1 and mGluR5 are known to contribute to the induction of mGluR-LTD at Sc-CA1 synapses. Both receptors are expressed in the hippocampus, both couple to Gα_q/11_ to signal to the canonical phospholipase C (PLC)/IP3 pathway and have been also proposed to form heterodimers (25). However, although the receptors have been generally regarded as interchangeable and able to compensate for each other’s activity, there is evidence that mGluR1 and mGluR5 can differ functionally. mGluR1 is involved in intracellular calcium release and the maintenance phase of mGluR-LTD, while mGluR5 is involved in potassium current influx and the induction phase of mGluR-LTD (26, 27).

**Figure 1:**
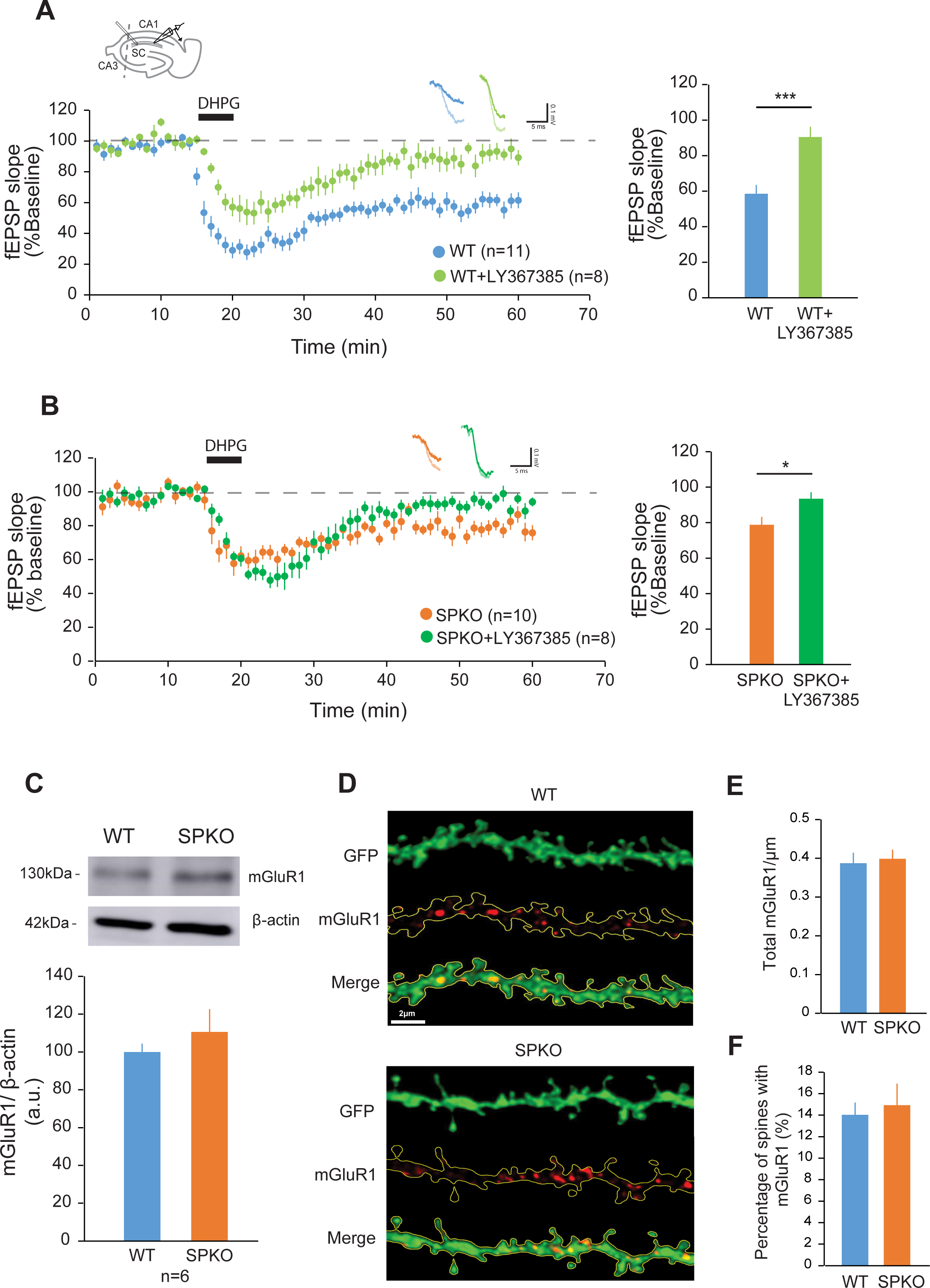
mGluR1 activity is necessary for mGluR-LTD in SPKO mice. (**A**) (*Left)* Time course of normalized fEPSP in WT (n=11 slices, N=9 mice) and WT+LY367385 (n=8 slices, N=5 mice). Diagram showing the positions of stimulating and recording electrodes at the Sc-CA1 area is included at the top. (*Right)* Quantification of average mGluR-LTD in the last 10 min of the recording. Mean ± SEM: WT=59.1%±4.8, WT+LY367385=90.5%±5.7, p<0.001 (Mann-Whitney test). (**B**) (*Left)* Time course of normalized fEPSP in SPKO (n=10 slices, N=7 mice) and SPKO+LY367385 (n=8 slices, N=4 mice). (*Right)* Quantification of average mGluR-LTD in the last 10 min of the recording. Mean ± SEM: SPKO=76.5%±4.4, SPKO+LY367385= 93.4%±3.7, p<0.05 (Mann-Whitney test). (**C)** Immunoblot of total mGluR1 protein from hippocampi of WT (n=6) and SPKO (n=6) mice. Mean ± SEM: WT=100.0%±4.4, SPKO=110.6%±12.0, p>0.05 (two-tailed unpaired t test). (**D)** Representative image of CA1 pyramidal neuron tertiary dendrites immunostained with mGluR1 antibody in WT (n=10) and SPKO (n=10) acute hippocampal slices. Scale bar=2 μm. (**E)** Quantification of mGluR1 puncta density in WT and SPKO tertiary dendrites. Mean ± SEM: WT=0.39±0.03, SPKO=0.40±0.02, p>0.05 (two-tailed unpaired t test). (**F)** Quantification of percentage of dendritic spines containing mGluR1 puncta in WT and SPKO tertiary dendrites. Mean ± SEM: WT=14.0%±1.1, SPKO=14.9%±2.0, p>0.05 (two-tailed unpaired t test).

To begin exploring the mechanisms underlying weakened mGluR-LTD in the absence of SP, we examined the individual contribution of mGluR1 and mGluR5 to the plasticity. To assess mGluR1 activity, we bath-applied the selective mGluR1 antagonist LY-367385 (100 µM) during the recording of LTD induced by DHPG. In WT, inhibition of mGluR1 significantly decreased the magnitude of the depression and blocked mGluR-LTD expression (90.5±5.7% of baseline, n=8; **Fig. 1A**). In SPKO mice, despite mGluR-LTD being already strongly reduced (76.5±4.4% of baseline, **Fig. 1B**), application of LY-367385 further decreased and fully blocked LTD (93.4±3.7% of baseline, n=8; **Fig. 1B**). To test whether the deficit in mGluR-LTD in SPKO mice could be due to reduced expression or mis-localization of mGluR1, we examined mGluR1 protein abundance and dendritic localization in the SPKO hippocampus. We found that both the total abundance of mGluR1 determined by Western blot (WT 100.0±4.4%, SPKO 110.6±12.0, n=6 mice per group; **Fig. 1C**) and the density of immunolabeled mGluR1 puncta in dendrites of CA1 pyramidal neuron (WT 0.39±0.03/µm n=10, SPKO 0.40±0.02/µm n=10; **Fig. 1D, E**) were not significantly altered in absence of SP. Moreover, in both WT and SPKO mice, ∼15% of dendritic spines in CA1 pyramidal neuron similarly displayed mGluR1 expression (**Fig. 1F**). Collectively, these data show that the lack of SP does not alter mGluR1 expression, localization and signalling in the hippocampus and demonstrate that mGluR1 activity is essential to support the residual mGluR-LTD in SPKO mice.

mGluR5 is abundantly expressed in the hippocampus and was shown to induce mGluR-LTD (27, 28). To test the contribution of mGluR5 activity to mGluR-LTD in absence of SP, the mGluR5-specific antagonist 2-methyl-6-(phenylethynyl) pyridine (MPEP, 10 μM) was applied to hippocampal slices during mGluR-LTD recording. As previously shown, in WT mice the DHPG-induced LTD was reduced in the presence of MPEP although not blocked (78.0±5.0% of baseline, n=10; **Fig. 2A**). Surprisingly, we found that inhibition of mGluR5 activity did not alter the magnitude of mGluR-LTD in SPKO mice (75.2±6.7% of baseline, n=9; **Fig 2B**). It is worth noting that inhibition of mGluR5 in WT (78.0±5.0%) results in LTD of magnitude similar to control untreated SPKO mice (76.5±4.4%). Since these findings suggested a potential impairment in mGluR5 activity in absence of SP, we next examined total mGluR5 protein expression by Western blot and found no significant difference in WT and SPKO hippocampus (**Fig. 2C**). Similarly, visualization of mGluR5 by immunolabeling of WT and SPKO hippocampal slices did not reveal detectable differences in the density of total (intracellular and surface) mGluR5-positive puncta in CA1 dendrites (**Fig. 2D, E**); congruent with this, the fraction of spines containing total mGluR5 was similar between genotypes (**Fig. 2F**). Altogether, these data indicate that although total mGluR5 expression remains normal in SPKO mice, mGluR5 function is impaired and is no longer required for mGluR-LTD in absence of SP.

**Figure 2:**
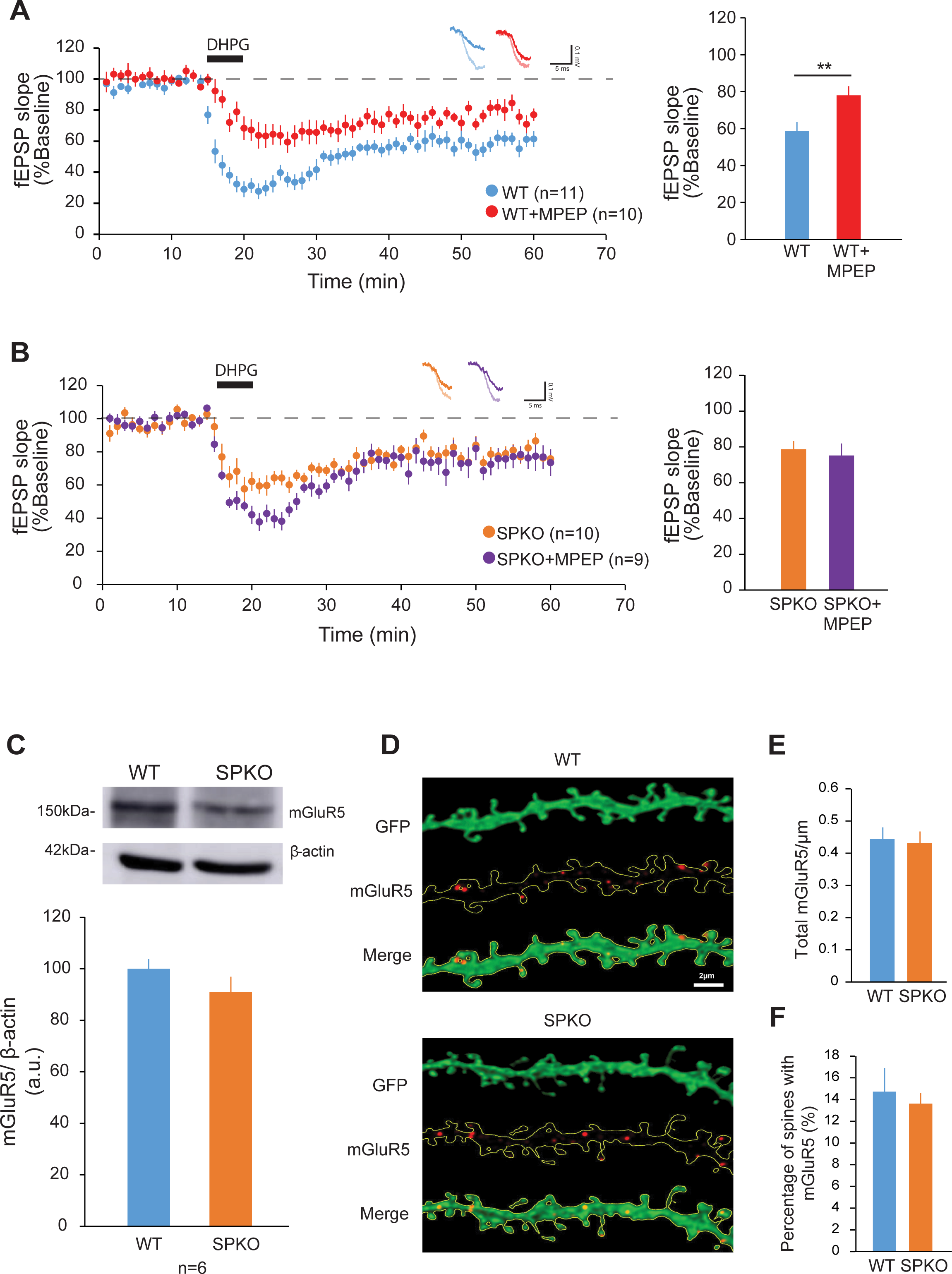
mGluR-LTD in SPKO mice is mGluR5 activity-independent. (**A)** (*Left)*Time course of normalized fEPSP in WT (n=11 slices, N=9 mice) and WT+MPEP (n=10 slices, N=7 mice). (*Right)* Quantification of average mGluR-LTD in the last 10 min of the recording. Mean ± SEM: WT=59.1%±4.8, WT+MPEP=78.0%±5.0, p<0.001 (Mann-Whitney test). (**B)** (*Left)*Time course of normalized fEPSP in SPKO (n=10 slices, N=7 mice) and SPKO+MPEP (n=9 slices, N=4 mice). (*Right)* Quantification of average mGluR-LTD in the last 10 min of the recording. Mean ± SEM: SPKO=76.5%±4.4, SPKO+MPEP=75.2%±6.7, p>0.05 (Mann-Whitney test). (**C)** Immunoblot of total mGluR5 protein from hippocampi of WT (n=6) and SPKO (n=6) mice. Mean ± SEM: WT=100.0%±3.7, SPKO=91.0%±5.9, p>0.05 (two-tailed unpaired t test). (**D)** Representative image of CA1 pyramidal neuron tertiary dendrites immunostained with mGluR5 antibody on WT (n= 14) and SPKO (n=11) acute hippocampal slices. Scale bar=2 μm. (**E)** Quantification of mGluR5 puncta density in WT and SPKO tertiary dendrites. Mean ± SEM: WT=0.44±0.04, SPKO=0.43±0.04, p>0.05 (two-tailed unpaired t test). (**F)** Quantification of percentage of dendritic spines containing mGluR5 puncta in WT and SPKO tertiary dendrites. Mean ± SEM: WT=14.7%±2.2, SPKO=13.6%±1.0, p>0.05 (two-tailed unpaired t test).

### mGluR5 surface level is significantly decreased in absence of synaptopodin

We reasoned that despite its normal abundance, mGluR5 expression at the neuronal surface and/or coupling to downstream signalling pathways could be affected by the lack of SP. Thus, to measure the surface level of mGluR5 and mGluR1, we immunolabeled intact (non-permeabilized) hippocampal slices with antibodies which specifically bind the extracellular domain of either receptor. We found that the mean fluorescence intensity of mGluR5 surface puncta was significantly reduced in the *stratum radiatum* of area CA1 in SPKO (6.8±0.7) compared to WT mice (17.5±1.4) **(Fig. 3A, B)**, whereas mGluR1 fluorescence intensity was unchanged (WT: 6.5±0.8, SPKO: 5.8±0.6) (**Fig. 3C, D**). The decrease in surface mGluR5 in SPKO mice occurred specifically in pyramidal neurons and not interneurons or glial cells, as indicated by reduced density of mGluR5 surface puncta along dendrites of GFP-labeled pyramidal neuron in the *stratum radiatum* of area CA1 (**Fig. 3E, F**).

**Figure 3:**
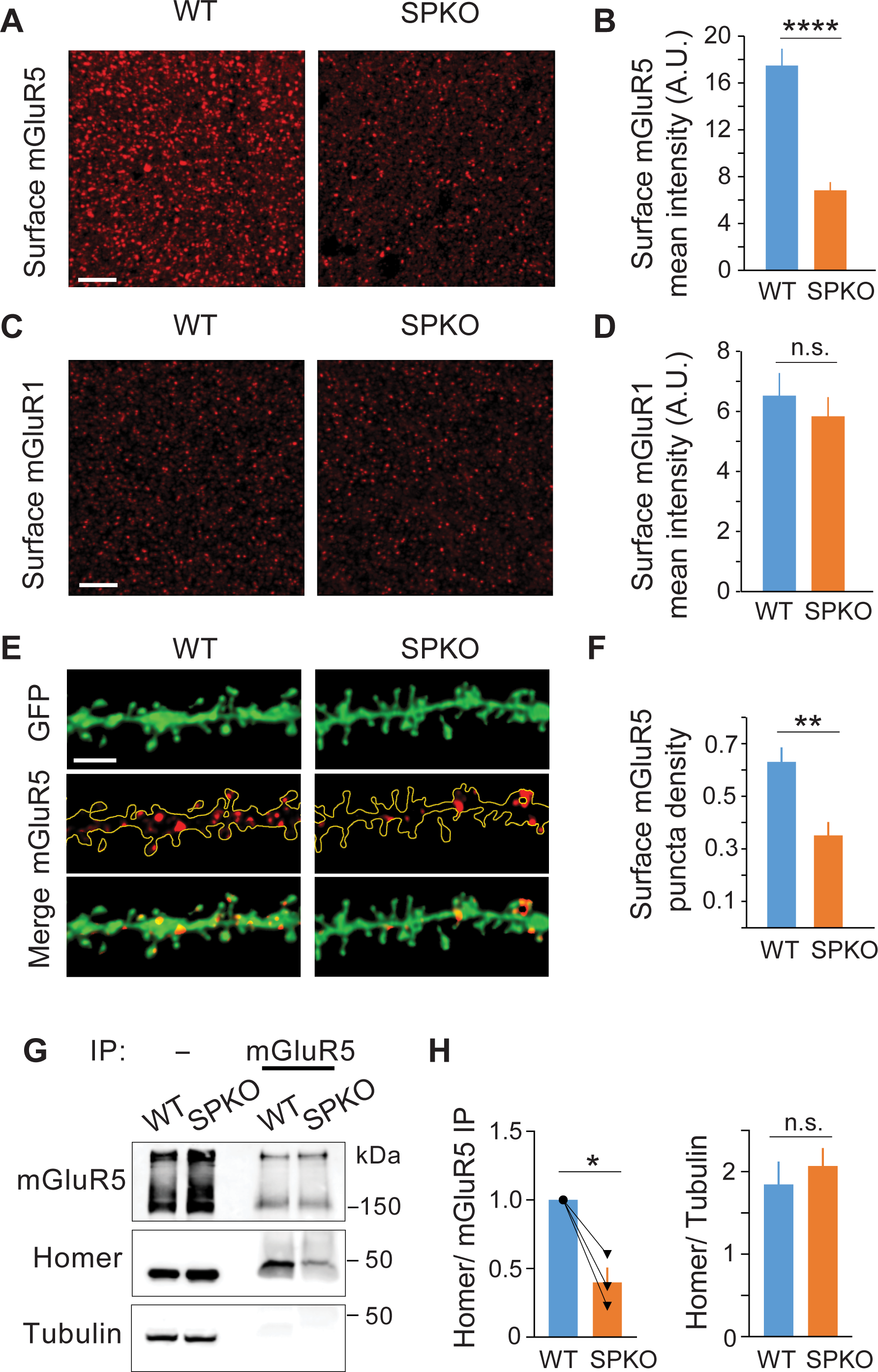
mGluR5 surface level on CA1 pyramidal neurons is significantly decreased and interaction with Homer is impaired in SPKO mice. (**A**) Representative images of surface mGluR5 in CA1 *stratum radiatum* of WT (n=10) and SPKO (n=9) mice. Scale bar=15 μm. (**B**) Quantification of mGluR5 puncta average intensity from images as in (A). Mean ± SEM: WT=17.5±1.4, SPKO=6.8±0.7, p<0.001 (two-tailed unpaired t test). (**C**) Representative images of surface mGluR1 in CA1 *stratum radiatum* of WT (n=5) and SPKO (n=8) mice. Scale bar=15 μm. (**D**) Quantification of mGluR1 puncta average intensity from images as in (C). Mean ± SEM: WT=6.5±0.8, SPKO=5.8±0.6, p>0.05 (two-tailed unpaired t test). (**E**) Representative images of CA1 pyramidal neuron tertiary dendrites immunostained with surface mGluR5 antibody in WT (n=11) and SPKO (n=11) acute hippocampal slices. Scale bar=3 μm. (**F**) Quantification of surface mGluR5 puncta density in CA1 pyramidal neuron tertiary dendrites. Mean ± SEM: WT=0.63±0.06, SPKO=0.35±0.05, p<0.01 (two-tailed unpaired t test). (**G)** Representative immunoblot of Homer co-immunoprecipitation (IP) with mGluR5 and expression in cortical tissue of WT and SPKO mice**. (H)** Quantification of Homer co-precipitated with mGluR5 and total abundance in WT and SPKO mice (N=3). Co-precipitated Homer: mean ± SEM SPKO/WT ratio 0.40±0.11, p=0.032 (paired t-test). Total Homer: mean ± SEM WT 1.8±0.28, SPKO 2.1±0.22, p=0.563 (unpaired t-test).

The clustering and mobility of mGluR5 at the neuronal surface is dependent on interaction with the scaffold protein Homer (29), which also physically connects receptors present at the plasma membrane with intracellular IP3R and RyR located in the ER (30–32). It is well established that absence of SP causes loss of the spine apparatus and therefore loss of stable ER within spines (18). We reasoned that lack of recruitment of Homer proteins to stable spine ER in SPKO mice might preclude local interaction with mGluR5. To explore this possibility, we immunoprecipitated mGluR5 from brain lysates of WT and SPKO mice and assessed co-precipitation of Homer (long form) by Western blot. As expected, Homer readily co-precipitated with mGluR5 in the WT; in contrast, Homer co-precipitation with the receptor was significantly reduced in SPKO mice (**Fig. 3G, H**) in absence of changes in total Homer abundance. Altogether, these results suggest that the lack of SP decreases the surface abundance of mGluR5 in pyramidal neurons *via* mechanism(s) involving impairment of the mGluR5-Homer association that normally promotes surface clustering of the receptors. We propose that deficits in mGluR5 surface expression and coupling to Homer underlie, in whole or in part, the failure of mGluR5 to contribute to mGluR-LTD in SPKO mice.

### mGluR-LTD is protein synthesis-independent in absence of synaptopodin

In juvenile and adult hippocampus, mGluR-LTD normally relies on rapid local translation of a specific group of proteins, termed “LTD proteins”, that are involved in the endocytosis and trafficking of AMPARs (11, 33). Blocking protein synthesis or eliminating specific “LTD proteins” results in severe impairment or abolishment of mGluR-LTD (11, 34). Protein synthesis can be directly induced by activation of group 1 mGluRs (35) and polyribosomes that are present in proximity to the spine apparatus (36, 37) that was also proposed to function as satellite secretory station (38, 39). Since the spine apparatus is absent in mice that lack SP, we tested whether *de novo* protein synthesis could be impaired in SPKO mice. Stimulation of protein synthesis is regulated by the extracellular signal-regulated kinases (ERK1/2) and the mechanistic target of rapamycin (mTOR) pathways that are both activated by group 1 mGluRs (33). Thus we used western blot assays with lysates of hippocampal slices treated with or without DHPG (100 μM, 5 min) to examine activation of the ERK1/2 and mTOR pathways. DHPG significantly increased ERK1/2 phosphorylation at Thr202/Tyr204 in WT, congruent with previous reports that the ERK1/2 pathway is activated and required for mGluR-LTD (**Fig. 4A**). In SPKO, DHPG induced ERK1/2 phosphorylation at a level comparable to WT (**Fig. 4A**) indicating that activation of the

**Figure 4:**
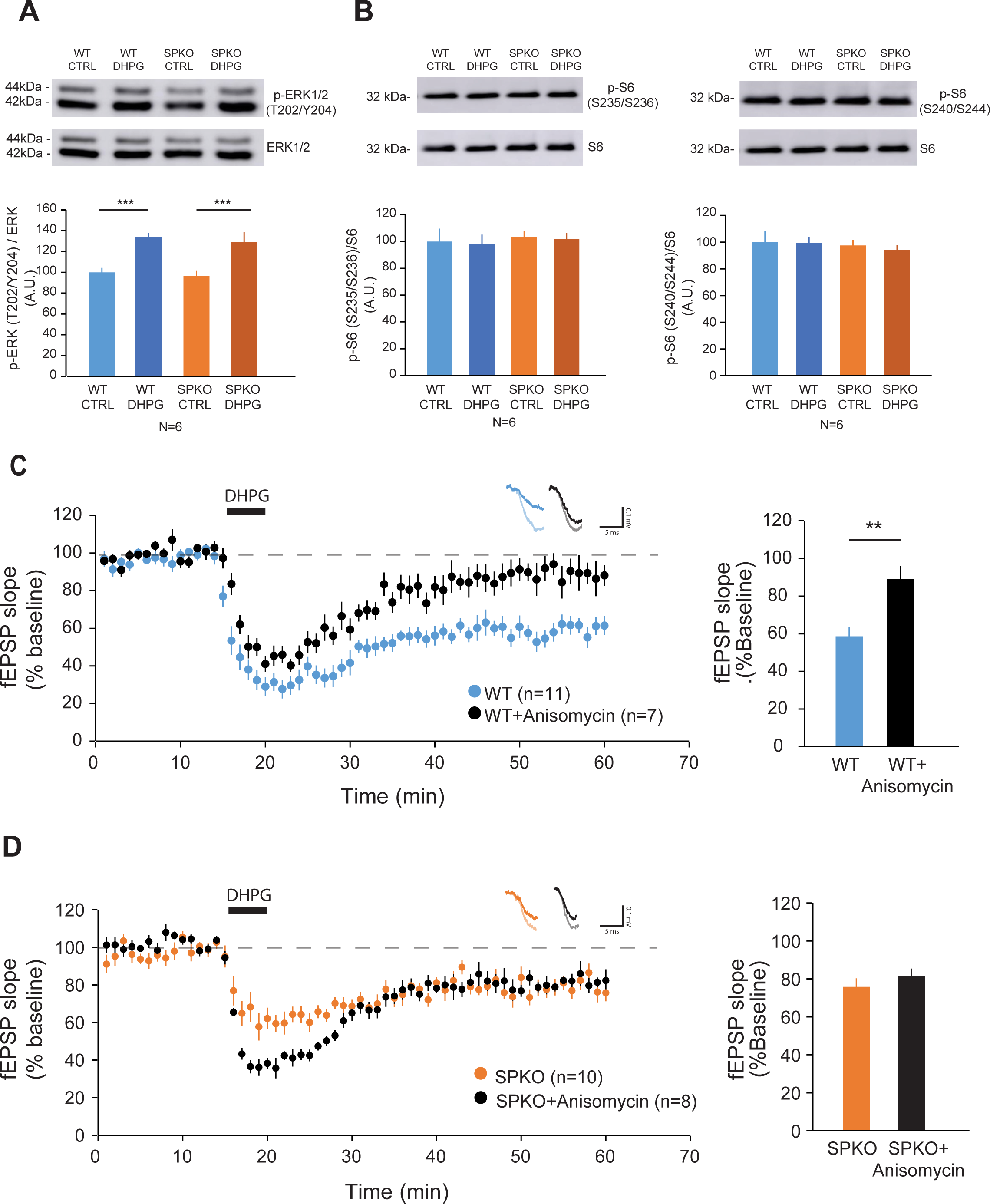
Hippocampal mGluR-LTD is protein synthesis independent in SPKO mice. (**A)** Representative immunoblot and quantification of phosphorylated ERK1/2 normalized to total ERK1/2 in WT and SPKO slices with or without DHPG treatment (100 μM, 5 min). Mean ± SEM: WT Ctrl=100.0%±4.3, WT+DHPG=134.2%±3.5, SPKO Ctrl =96.6%±4.8, SPKO+DHPG= 129.1%±9.4, p<0.01 (two-way ANOVA with Sidak’s test). (**B)** Representative immunoblot and quantification of (*left*) S6 phosphorylated at S235/S236 and (*right*) at S240/S244 normalized to total S6 in WT and SPKO with or without DHPG treatment (100 μM, 5min). Mean ± SEM p-S6 (S235/S236): WT Ctrl=100.0%±9.5, WT+DHPG=98.2%±6.9, SPKO Ctrl=103.4%±4.4, SPKO+ DHPG=101.8%±4.8, p>0.05 (two-way ANOVA with Sidak’s test). Mean ± SEM p-S6 (S240/S244): WT Ctrl=100.0%±8.0, WT+DHPG=99.4%±4.5, SPKO Ctrl=97.5%±4.2, SPKO+DHPG=94.3%±3.5, p>0.05 (two-way ANOVA with Sidak’s test). (**C)** (*Left)*Time course of normalized fEPSP in WT (n=11 slices, N=9 mice) and WT+Anisomycin (n=7 slices, N=3 mice). (*Right)* Quantification of average mGluR-LTD in the last 10 min of the recording. Mean ± SEM: WT=59.1%±4.8, WT+Anisomycin=89.0%±7.1, p<0.001 (Mann-Whitney test). (**D)** (*Left)*Time course of normalized fEPSP in SPKO (n=10 slices, N=7 mice) and SPKO+Anisomycin (n=8 slices, N=4 mice). (*Right)* Quantification of average mGluR-LTD in the last 10 min of the recording. Mean ± SEM: SPKO=76.5%±4.4, SPKO+Anisomycin=81.5%±3.9, p<0.05 (Mann-Whitney test).

ERK1/2 pathway by DHPG is not affected in absence of SP. To assess the mTOR pathway, we measured phosphorylation of the ribosomal protein S6 (Ser235/236, Ser240/244), a downstream target of mTORC1 signaling. We found that DHPG treatment did not affect S6 phosphorylation in either WT or SPKO, suggesting that group 1 mGluR signalling to mTORC1 is unchanged (**Fig. 4B**). The ERK1/2 and mTOR pathways are signalling cascades that regulate protein synthesis but their activities do not directly reflect actual *de novo* production of proteins. For a direct assessment of how the lack of SP might affect *de novo* protein synthesis during mGluR-LTD, we applied DHPG to hippocampal slices while perfusing anisomycin, a protein translation inhibitor, and recorded the fEPSPs. In WT, treatment with anisomycin blocked mGluR-LTD expression (89.0±7.1% of baseline; **Fig. 4C**), in line with reports that mGluR-LTD heavily relies on *de novo* protein synthesis

(11). Since protein synthesis is required for mGluR-LTD, we anticipated that the residual mGluR-LTD in SPKO mice would also be blocked by anisomycin. To our surprise, in SPKO mice anisomycin did not block mGluR-LTD and in fact, the treatment had no effect on the fEPSPs compared to vehicle control (81.5±3.9% of baseline; **Fig. 4D**) indicating that the residual mGluR-LTD in SPKO mice is protein synthesis-independent.

### In absence of synaptopodin mGluR-LTD is mediated *via* an endocannabinoid-dependent mechanism

The mechanisms of expression of mGluR-LTD at Sc-CA1 synapses undergo a developmental switch (40). Whereas mGluR-LTD in the juvenile/adult hippocampus is mediated by postsynaptic induction of protein synthesis and production of “LTD” proteins promoting removal of synaptic AMPARs (33), in neonatal animals mGluR-LTD is protein synthesis-independent and expressed through a decrease in presynaptic function (40, 41). As the residual mGluR-LTD in SPKO mice becomes protein synthesis-independent, a critical question to address is what mechanism(s) underlie functional synaptic weakening. In many brain regions, including hippocampus, endocannabinoid signalling can decrease neurotransmitter release at excitatory synapses and induce LTD (10). Endocannabinoids are produced post-synaptically in response to group 1 mGluR activation and act as retrograde signal back to the presynaptic terminal by binding to cannabinoid receptor type 1 (CB1) thereby causing synaptic depression (10, 42). To investigate whether residual mGluR-LTD in absence of SP might be produced by a presynaptic mechanism through endocannabinoid signaling, we tested the effect of the CB1 receptor inhibitor AM-251 on mGluR-LTD in SPKO mice. Surprisingly, inhibition of the CB1 receptor in SPKO hippocampal slices blocked mGluR-LTD induced by DHPG (91.9±4.2% of baseline; **Fig. 5A**). In contrast, application of AM-251 to WT slices did not significantly alter the magnitude of DHPG-induced LTD (66.7±3.9% of baseline; **Fig. 5B**). Together, these findings support the notion that mGluR-LTD is normally expressed post-synaptically but loss of SP causes a switch in the locus of plasticity to the presynaptic side.

**Figure 5:**
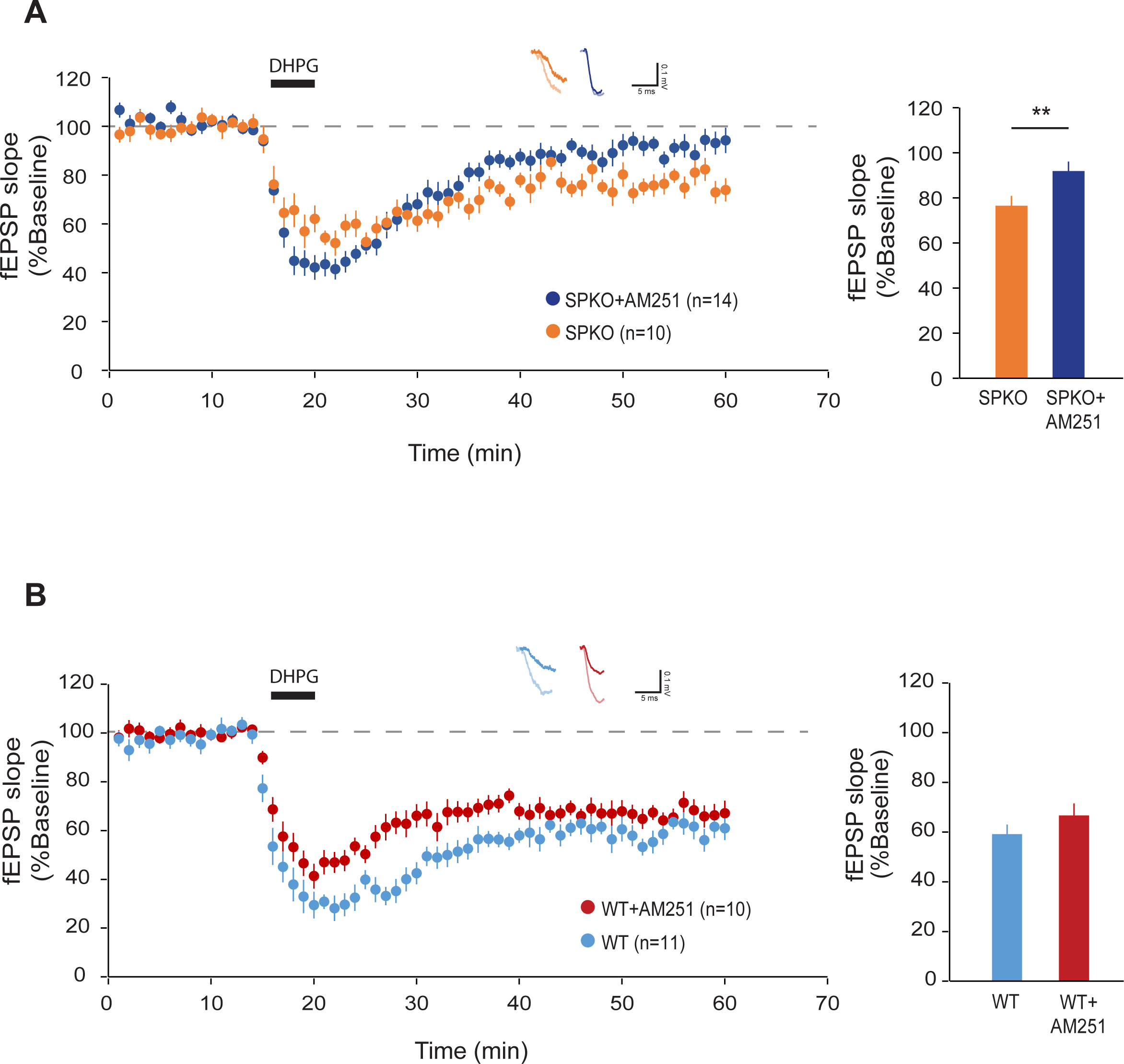
Residual mGluR-LTD in SPKO mice is mediated through endocannabinoid signaling. **(A)** (*Left*) Time course of normalized fEPSP in SPKO (n=10 slices, N=7 mice) and SPKO+AM251 (n=14 slices, N=6 mice). *(Right)* Quantification of average mGluR-LTD in the last 10 min of the recording. Mean ± SEM: SPKO=76.5%±4.4, SPKO+AM251= 91.9%±4.2, p<0.01 (Mann-Whitney test). **(B)** (*Left)*Time course of normalized fEPSP in WT (n=11 slices, N=9 mice) and WT+AM251 (n=10 slices, N=3 mice). (*Right)* Quantification of average mGluR-LTD in the last 10 min of the recording. Mean ± SEM: WT=59.1%±4.8, WT+AM251=66.7%±3.9, p>0.05 (Mann-Whitney test).

## Discussion

DHPG-induced LTD at Sc-CA1 synapses is strongly reduced in absence of SP, an actin-binding protein enriched in a subset of dendritic spines (24). In this study we identify alterations that occur at Sc-CA1 glutamatergic synapses in absence of SP and provide evidence of its crucial role in enabling postsynaptic expression and protein synthesis dependence of mGluR-LTD. We show that activation of mGluR1 is required for both robust LTD in WT and residual LTD in SPKO mice, whereas mGluR5 no longer contributes to LTD in absence of SP. In SPKO mice, mGluR5 surface expression is selectively reduced in CA1 pyramidal neurons and its association with Homer proteins impaired. We further find that inhibition of *de novo* protein synthesis which blocks mGluR-LTD in the WT has no effect on mGluR-LTD in absence of SP. Instead, in SPKO mice mGluR-LTD is blocked by inhibition of the CB1 receptor, which is activated by endocannabinoids produced downstream of group 1 mGluR signalling. Overall, our results point to a key role of SP in enabling postsynaptic expression of mGluR-LTD.

It was previously reported that the concomitant inhibition of mGluR1 and mGluR5 is required to block induction of mGluR-LTD at Sc-CA1 synapses and that inhibition of mGluR1, but not mGluR5, can block LTD expression (27). Our results confirm previous work and further show that inhibition of mGluR1 blocks both mGluR-LTD in WT and the residual LTD observed in SPKO mice, indicating that in absence of SP mGluR1 remains capable of promoting synaptic weakening. This conclusion is congruent with the observation that global and surface expression of mGluR1 is not altered in SPKO mice. In contrast, mGluR5 function in mGluR-LTD is lost in absence of SP, an impairment potentially caused, in whole or in part, by the reduced surface expression of the receptor in CA1 pyramidal neurons. Deficits in mGluR5 surface expression could be linked to its reduced association with Homer scaffolds in absence of SP. The long variants of Homer proteins form tetrameric hubs (43, 44) that crosslink postsynaptic surface mGluR1a/mGluR5 with intracellular IP3R and RyR in ER membranes facilitating Ca^2+^ signalling (30, 32). Association with Homer proteins also modifies the lateral mobility of mGluR5 at the neuronal surface by promoting receptor clustering and countering activity-induced lateral diffusion (29). Notably, the interaction of mGluR5 with Homer was shown to be necessary for the ability of the receptor to induce mGluR-LTD (45). In absence of SP, the spine apparatus - a source of stable ER within spines - does not form (18), affecting the dendritic localization of ER-bound IP3R and RyR and likely the consequent recruitment of Homer scaffolds to spine ER. In addition, a putative Homer-binding motif was detected in SP isoforms suggestive of a potential direct interaction between the proteins (15). Overall, our results are consistent with a model whereby reduced mGluR5-Homer crosslinking and mGluR5 surface clusters in absence of SP compromise the capacity of the receptor to contribute to mGluR-LTD in SPKO mice. It is important to note that unlike mGluR5, mGluR1 is encoded by multiple splice variants that lack the Homer-binding motif (46, 47), with the exception of the mGluR1a variant that however is mostly expressed in interneurons in area CA1 (48). It is likely that alternative Homer-independent mechanisms, which appear to remain functional in absence of SP, contribute to regulate mGluR1 surface expression and stability at excitatory synapses (49).

mGluR-LTD has been shown to heavily rely on *de novo* protein synthesis, which is regulated by signalling to the ERK and the mTOR pathways. ERK1/2 activation was shown to be required for mGluR-LTD and several studies have demonstrated that mGluR5 is responsible for ERK activation (50, 51). Here we show that ERK1/2 phosphorylation is significantly increased after DHPG stimulation in WT, in agreement with others, and that ERK1/2 is equally activated in SPKO mice. Ronesi *et al.* have previously shown that the disruption of mGluR5-Homer interaction in hippocampus impairs mGluR-LTD but does not affect ERK1/2 activation, similarly to what we observe in SPKO mice in which mGluR5 association with Homer is impaired. Notably, in knock-in mice with a mutation in mGluR5 that abolishes its binding to Homer, mGluR-LTD becomes protein synthesis-independent (52) similarly to SPKO mice.

Given the more abundant expression of mGluR5 in the hippocampus compared to mGluR1 it might be counterintuitive that mGluR1 activity is the predominant participant in mGluR-LTD at Sc-CA1 synapses. However, multiple independent reports converge to indicate that mGluR1 is critical to the regulation of synaptic weakening and synapse elimination in the CNS through both post-and presynaptic mechanisms (27, 53–58). At hippocampal synapses, in addition to functional synaptic weakening, mGluR1 is critically involved in the elimination of dendritic spines during mGluR-LTD supporting the idea that mGluR1, although sparse, plays an indispensable role in hippocampal mGluR-LTD (24). In absence of SP, mGluR1 capacity to promote spine loss is impaired (24) whereas here we report that its contribution to synaptic weakening appears to remain intact. We reason that the structural and the functional changes induced by the LTD are not coupled to each other and are instead mediated via distinct pathways, meaning that the loss of structural plasticity does not necessarily translate to the loss of functional plasticity and vice versa. A potential scenario is that mGluR1-induced spine loss relies on postsynaptic effector mechanisms (*e.g.,* protein synthesis) that are lost in absence of SP whereas presynaptic expression of synaptic weakening driven by mGluR1 is preserved. Although presynaptic expression of hippocampal mGluR-LTD was mostly recorded in neonatal animal (≤2 week-old), there is also evidence that a presynaptic component contributes to the plasticity at later ages (7, 59). In absence of SP, the ensuing postsynaptic deficits may unmask existing presynaptic changes during the LTD that might be further strengthened as compensatory mechanism. Nevertheless, our results reinforce the notion that mGluR1 and mGluR5, though traditionally considered to share the same molecular pathways, are functionally different from each other.

Dendritic spines are functionally and structurally heterogeneous. The presence of SP in a subset of mushroom spines is critical for spine stabilization (60) and for the generation of stronger postsynaptic responses compared to spines lacking SP (20, 21). Spines harbouring SP and a spine apparatus support synaptic plasticity, as demonstrated by the findings that SPKO mice show impaired LTP (18, 22, 61) and mGluR-LTD (24). The locus of expression for mGluR-LTD can be influenced by various factors including hippocampus development, aging, subregion and experimental protocol (40, 62–64). In this study, we show for the first time that SP, a postsynaptic protein, controls the locus of expression of mGluR-LTD. The loss of SP eliminates the *de novo* protein synthesis dependency of hippocampal mGluR-LTD that occurs at mature synapses and switches the system to rely on the endocannabinoid signalling that produces LTD through presynaptic mechanism as in neonatal animals (40). It is important to note that expression of SP and consequently the formation of the spine apparatus are developmentally regulated. SP is gradually expressed in the hippocampus during the postnatal period, starting postnatally around the first week and is strongly expressed during adult life (65, 66) parallel to the switch of mGluR-LTD from presynaptic to postsynaptic, protein synthesis-dependent mechanisms of expression.

In Fragile X syndrome (FXS), a neurodevelopmental disorder characterized by enhanced mGluR-LTD, basal protein synthesis is increased and SP is found more abundantly in dendritic spines (67, 68). The fragile X messenger ribonucleoprotein (FMRP) - silenced in FXS – regulates RNA translation and was found associated with the spine apparatus (69). We propose that stable SP-positive spines containing the spine apparatus provide the postsynaptic locus for mGluR-LTD and alterations in SP expression and formation/stability of the spine apparatus may contribute to abnormal synaptic plasticity in FXS and other neurodevelopmental disorders marked by altered mGluR-LTD.

## Materials and Methods

### Animals

All procedures were according to protocols approved by the guidelines of the Canadian Council on Animal Care and the McGill University Comparative Medicine and Animal Resources animal handling protocols 5057. C57Bl6 and L15 transgenic mice expressing membrane-targeted MARCKS (myristoylated alanine-rich protein kinase C substrate)–enhanced GFP under the Thy-1 promoter in a subpopulation of CA1 cells were used as WT (70). SPKO and L15S were used as SPKO and were previously described(18, 19). Mice were fed *ad libitum* and housed with a 12 h light/dark cycle. WT and SPKO male mice were used for experiments.

### Drugs and chemicals

S-3,5-Dihydroxyphenylglycine (DHPG), 2-methyl-6-(phenylethynyl) pyridine hydrochloride (MPEP), (S)-(+)-α-Amino-4-carboxy-2-methylbenzeneacetic acid (LY367385), anisomycin and N-(Piperidin-1-yl)-5-(4-iodophenyl)-1-(2,4-dichlorophenyl)-4-methyl-1H-pyrazole-3-carboxamide (AM-251) were purchased from Tocris Bioscience. DHPG and MPEP were prepared in ddH_2_O. LY367385 was prepared in 1.1eq NaOH solution. AM-251 and anisomycin were prepared in DMSO.

### Electrophysiology

Hippocampal slices were obtained from P30-P40 old WT or SPKO mice. Mice were deeply anesthetized with isoflurane and killed by decapitation. Slices (400 µm) were cut on a vibratome (Leica Microsystems, VT1200S) in a sucrose-based solution containing the following (in mM): 280 sucrose, 26 NaHCO_3_, 10 glucose, 1.3 KCl, 1 CaCl_2_, and 10 MgCl_2_ and were transferred at 32°C in regular ACSF containing the following (in mM): 124 NaCl, 5 KCl, 1.25 NaH_2_PO_4_, 2 MgSO_4_, 26 NaHCO_3_, 2 CaCl_2_, and 10 glucose saturated with 95% O_2_/5% CO_2_ (pH 7.3, 300 mOsm) for 15 min before resting at room temperature (RT) for 1 h in oxygenated (95% O_2_/5% CO_2_) ACSF. To assess mGluR-LTD, slices were placed into a heated (32°C) recording chamber of an upright microscope (DM LFSA Microsystems) and perfused continuously with oxygenated ACSF or ACSF containing indicated drugs. S-DHPG (100 μM) was applied for 5 minutes. Field excitatory postsynaptic potentials (fEPSPs) were recorded in the *stratum radiatum* of the CA1 region by using glass microelectrodes filled with 3 M NaCl. GABA_A_ receptor-mediated inhibition was blocked with 100 μM picrotoxin, and the area CA1 was surgically isolated from CA3 to avoid epileptiform activity. fEPSPs were elicited at 0.1 Hz by a digital stimulator that fed by a stimulation isolator unit. All data analyses were performed with custom-written software in Igor Pro 8 (Wavemetrics). fEPSP slope was measured as an index of synaptic strength.

### Immunohistochemistry

Hippocampal slices of 100 µm were obtained from P30 to P40 old L15S or L15 mice. The slices were incubated in ACSF at RT for 1 h for recovery and fixed in 0.1 M Phosphate buffer (PB) containing 4% PFA, pH 7.4, overnight at 4°C. After fixation, slices were washed in 0.1 M PB, permeabilized in 0.4% Triton X-100, and blocked with 1.5% heat-inactivated horse serum overnight at 4°C. Slices were incubated with primary anti-mGluR1 antibody (1:500, BD transduction, #610965) and primary anti-mGluR5 antibody (1:250, Millipore, #3352426) in permeabilizing solution for 3 days at 4°C, washed with PB, and incubated with anti-rabbit secondary antibody conjugated to DyLight 649 (1:500; Jackson ImmunoResearch Laboratories) for 3 hours, washed, and mounted with DAKO Fluorescent Mounting medium (Dako Canada) onto microscope slides before imaging and subsequent blinded analysis. Spine images were taken in z stacks using a Leica Microsystems SP8 confocal microscope with oil-immersion 63x objective at 6x zoom-in. The images were deconvolved and analyzed using software Huygens (Scientific Volume Imaging) and Imaris (Oxford Instruments) respectively.

For the detection of surface level of mGluR1 and mGluR5, the protocol was the same as above except that Triton X-100 was omitted. The slices were incubated with anti-mGluR1 (1:100, Alomone labs, CAT# AGC006) and anti-mGluR5 (1:100, Alomone labs, CAT# AGC007) that target the N-terminal extracellular tail. The images were taken at the *stratum radiatum* of CA1 hippocampus with oil immersion 63x objective at 0.75x zoom-in.

### Immunoblotting and drug treatment

Hippocampal slices (400 µm thick) were obtained from WT and SPKO mice at p30-40. The slices were incubated in ACSF at 32°C for 3 hours before treatment with S-DHPG (100 μM) for 5 min. After treatment, the slices were rapidly homogenized in ice-cold RIPA (150 mM NaCl, 1% Nonidet P-40, 0.5% sodium deoxycholate, 0.1% sodium dodecylsulfate, 50 mM Tris-HCl (pH 8.0) with protease and phosphatase inhibitors (1X Roche Complete Mini, 5 mM NaF, 1 mM sodium orthovandate, 1 mM PMSF). Homogenates were sonicated and centrifuged for 5 min at 13,200 rpm. The supernatant was extracted and mixed with 2x Laemmli sample buffer to produce loading samples. Samples were boiled, resolved on SDS-PAGE, transferred to nitrocellulose, and stained with primary antibodies overnight. The following antibodies were obtained from Cell Signaling Technology (Danvers, MA): ERK1/2 (1:2000), p-ERK1/2-Thr202/Tyr204 (1:2000), S6 (1:1000), p-S6-Ser235/236-Ser240/244 (1:1000) except anti-β-actin (Sigma-Aldrich; 1:10,000) used for loading control. mGluR1 (1:1000, BD transduction, #610965) and mGluR5 (1:1000, Millipore, #3352426) antibodies were blotted in control slices for WT and SPKO. Horseradish peroxidase (HRP)-conjugated secondary antibodies goat anti-rabbit (abcam; 1:5000) or goat anti-mouse (Bio-Rad Laboratories; 1:10,000) were applied for 1 h. The blots were imaged with Amersham Imager 600 and the band intensities were analyzed using the open-source program Fiji (71).

### Immunoprecipitation

Brain cortices of 1 month-old mice were manually homogenized on ice with Potter-Elvehjem tissue homogenizer and vortexing (∼ 60 min total, each stroke followed by 10 min incubation on ice) in a buffer of 20 mM Tris-HCl pH 7.4, 5 mM EDTA, 100 mM NaCl, 1% Triton X-100 (10 μl/mg tissue) supplemented with cocktails of protease and phosphatase inhibitors. The homogenized tissue was centrifuged at 20,500 x *g* for 20 min at 4°C and protein content in the supernatant measured by Bradford assay. Equal amount of protein (2 mg/IP) were incubated overnight at 4°C with rabbit anti-mGluR5 (7 μl/IP; Alomone Labs Cat# AGC-007, RRID:AB_2039991) bound to protein G-coupled magnetic beads (40 μl Dynabeads/sample; Invitrogen Waltham, MA). The beads were washed 2 times with homogenization buffer, 2 times with PB with 0.1% Triton X-100 and once with PB (10 min rotation at 4°C per wash). Bound proteins were eluted in denaturing sample buffer, heated at 95°C for 5 min and separated on 8% SDS-PAGE before transfer to nitrocellulose membrane and immunoblot with rabbit anti-pan-Homer (Santa Cruz Biotechnology Cat# sc-15321, RRID:AB_2120999; 1:250), rabbit anti-mGluR5 (Abcam Cat# ab76316, RRID:AB_1523944; 1/500), mouse anti-γ-Tubulin (Sigma-Aldrich Cat# T6557, RRID:AB_477584; 1:2500) followed by detection with HRP-conjugated secondary antibodies and chemiluminescent substrate (Immobilon Western; Millipore-Sigma, Burlington, MA). Bands were visualized with Azure 600 Imaging System (Azure Biosystems, Dublin, CA) and quantified with the program Fiji.

## Acknowledgements

This work was supported by National Institute of Mental Health R01MH108614 to A.F.; Canadian Institutes of Health Research MOP 86724 to R.A.M.; NSERC Discovery R.A.M. and the Norman Zavalkoff Family Foundation to R.A.M.; Richard and Edith Strauss Postdoctoral Fellowship in Medicine to Y.I. We thank members of the A.F. and R.A.M. laboratories for comments on the manuscript; and Francois Charron for excellent technical assistance. The authors declare no competing financial interests.

## Author contributions

Y.I., R.A.M., and A.F. designed research; P.Y.W., L.J., C.D.S. performed research; P.Y. W., L.J., C.D.S., Y.I. analyzed data; P.Y.W., Y.I, R.A.M. and A.F. wrote the paper.

