## Supplemental Figure 1 for "Loss of synaptopodin impairs mGluR5 and protein synthesis dependent mGluR-LTD at CA3-CA1 synapses"

Supplementary Figure 1

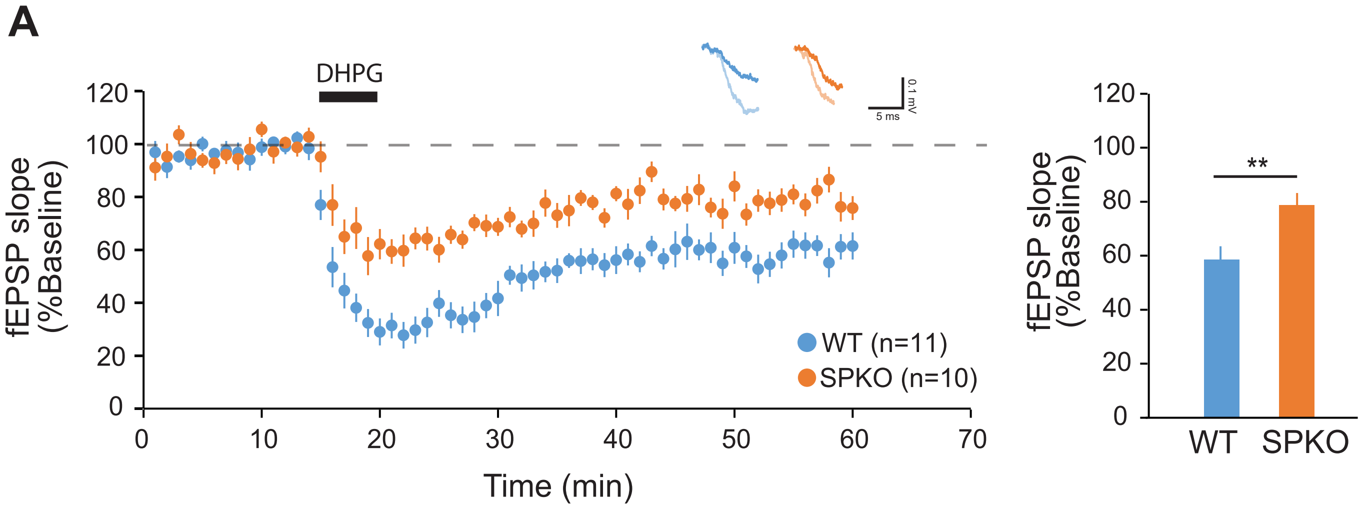

### Supplementary Figure legends

**Suppl. Figure 1: mGluR-LTD is significantly impaired in SPKO mice compared to WT**. (*Left)* Time course of normalized fEPSP in WT (n=11 slices, N=9 mice) and SPKO (n=10 slices, N=7 mice). (*Right)* Quantification of average mGluR-LTD in the last 10 min of the recording. Data are mean ± SEM. WT=59.1%±4.8, SPKO=76.5%±4.4. $p<0.01$ (Mann-Whitney test).
